# Effects of systemic oxytocin receptor activation and blockade on risky decision making in female and male rats

**DOI:** 10.1101/2024.05.13.593981

**Authors:** Mojdeh Faraji, Omar A. Viera-Resto, Brenden J. Berrios, Jennifer L. Bizon, Barry Setlow

**Affiliations:** Department of Psychiatry, University of Florida; Center for Addiction Research and Education, University of Florida; Department of Neuroscience, University of Florida; McKnight Brain Institute, University of Florida

**Keywords:** punishment, decision making, oxytocin, rat, risk taking

## Abstract

The neuropeptide oxytocin is traditionally known for its roles in parturition, lactation, and social behavior. Other data, however, show that oxytocin can modulate behaviors outside of these contexts, including drug self-administration and some aspects of cost-benefit decision making. Here we used a pharmacological approach to investigate the contributions of oxytocin signaling to decision making under risk of explicit punishment. Female and male Long-Evans rats were trained on a risky decision-making task in which they chose between a small, “safe” food reward and a large, “risky” food reward that was accompanied by varying probabilities of mild footshock. Once stable choice behavior emerged, rats were tested in the task following acute intraperitoneal injections of oxytocin or the oxytocin receptor antagonist L-368,899. Neither drug affected task performance in males. In females, however, both oxytocin and L-368,899 caused a dose-dependent reduction in preference for large risky reward. Control experiments showed that these effects could not be accounted for by alterations in food motivation or shock sensitivity. Together, these results reveal a sex-dependent effect of oxytocin signaling on risky decision making in rats.

## Introduction

Oxytocin (OT) is a neuropeptide synthesized primarily by magnocellular neurons in the paraventricular (PVN), supraoptic (SON), and accessory nuclei of the hypothalamus (Horn & Swanson, 2012). The axons of magnocellular neurons in PVN and SON project to the posterior pituitary and release OT from axon terminals into the bloodstream, where it plays pivotal roles in parturition and lactation via positive feedback signaling (Walter et al., 2021). In addition to acting as an endocrine hormone, however, OT is a central nervous system neurotransmitter involved in regulating social behavior, pair bonding, reproductive experience, and parental care. Axon projections from the PVN and SON nuclei deliver OT to the prefrontal cortex, hippocampus, amygdala, nucleus accumbens, thalamus, and other regions, all of which express the G-protein coupled oxytocin receptor (Jurek & Neumann, 2018; Knobloch et al., 2012).

The role of OT in the reproductive process goes beyond stimulating uterine contractions and lactation. Central OT signaling is reported to facilitate formation of pair bonding, and to induce maternal behavior and pup protection in rodents (Carcea et al., 2021; Pedersen et al., 1982; Pedersen & Boccia, 2003; Rickenbacher et al., 2017; Williams et al., 1994). OT signaling also modulates cognitive flexibility and anxiety- and depressive-like behaviors in female rats during the post-partum period (Albin-Brooks et al., 2017; Figueira et al., 2008; Wang et al., 2018). More broadly, OT is widely recognized to enhance prosocial behavior (Liu et al., 2019; Marsh et al., 2020). In humans, intranasal OT is reported to increase trust (Baumgartner et al., 2008; Kosfeld et al., 2005), improve identification of emotions expressed by photographed faces (Fischer-Shofty et al., 2010; Lischke et al., 2012), enhance empathy (specifically in participants worse at identifying other people’s emotions during conversation) (Bartz et al., 2010), and strengthen in-group cooperation (De Dreu, 2012).

While research on the neural circuits underlying central OT signaling is still ongoing, it is believed that OT affects behavior and cognition at least in part through interactions with reward and stress systems (King et al., 2020; Love, 2014; Takayanagi & Onaka, 2021). OT is additionally tightly connected to the neurobiology of mental health due to major contributions of reward processing dysfunctions to risks of psychiatric disorders (Pujara & Koenigs, 2014; Whitton et al., 2015). In fact, research in humans reveals altered peripheral OT levels in individuals with psychiatric disorders (Ferreira & Osório, 2022; Kirsch, 2015; Oh et al., 2018), and fMRI studies show differential activity patterns in patients compared to healthy controls when treated with intranasal oxytocin (Pincus et al., 2010; Wang et al., 2017). In the context of substance use disorders, intranasal OT attenuates tobacco and cannabis craving in humans (McRae-Clark et al., 2013; Miller et al., 2016) as well as drug-seeking behaviors in rodents (Ballas et al., 2021; Baracz et al., 2014; Baracz et al., 2012; Carson, Cornish, et al., 2010; Cox et al., 2013; Kovács et al., 1985; Leong et al., 2018; MacFadyen et al., 2016; Qi et al., 2009; Weber et al., 2018; Zhou et al., 2014).

Psychiatric disorders are associated with alterations in several forms of cost-benefit decision making, including risky decision making (Chen et al., 2020; Li et al., 2023; Pailing & Reniers, 2018; Reddy et al., 2014; Strawbridge et al., 2018). Research supports a role for OT signaling in risk-based decision making in social contexts; for example, intranasal OT reduces betting rate in men under social stress (Patel et al., 2015). There is, however, evidence that OT can also influence nonsocial decision making (Hansson et al., 2018; Kapetaniou et al., 2021; McRae-Clark et al., 2013; Michalopoulou et al., 2015; Miller et al., 2016; Plessow et al., 2021), including decision making involving risk. One study using the Iowa Gambling Task (IGT) found that intranasal OT administration reduced risk taking. In the IGT, participants choose between a high gain/high loss option (the “disadvantageous deck” that yields negative net earnings) and a low gain/low loss option (the “advantageous deck” that yields positive net earnings). In this study (Bozorgmehr et al., 2019), intranasal OT administration increased participants’ net earnings compared to control groups who received placebo or no treatments. In contrast, another study (Zebhauser et al., 2022) found that compared to control and placebo conditions, subjects given OT showed increased risk taking in the IGT. As these two studies employed nearly identical methods (same dose, route of administration, and general subject characteristics), the reason for their opposite outcomes is not clear, although it might be attributable to the use of a between-subjects (Bozorgmehr et al., 2019) vs. a within-subjects (Zebhauser et al., 2022) design. Another study assessed performance in patients with bulimia and healthy controls in the Balloon Analogue Risk Task, in which subjects pump up a virtual balloon to maximize earnings, without pumping so much as to pop the balloon (which results in loss of earnings). Acute OT administration had no main effect on risk taking in the subjects overall, but there was an interaction between the subject group and drug condition, such that compared to the placebo condition, participants in the healthy control group who received intranasal OT showed an increase in the number of “balloon explosions” (a measure of risk taking), while the patient group showed more risk-averse behavior in the OT condition (Leslie et al., 2019).

The results in humans on the effects of OT on risk-taking behavior, as well as the fact that these effects may vary across clinical populations, highlight a need for better understanding of the mechanisms by which OT exerts its effects on decision making. Animal models can be useful for gaining such mechanistic insights; to our knowledge, however, only one study to date has investigated the influence of OT on risk taking behavior. Using a “probability discounting task” in which rats choose between a small, guaranteed food reward and a large food reward delivered with varying probabilities (St Onge & Floresco, 2009), Tapp et al. found that intracerebroventricular OT administration dose-dependently reduced male rats’ preference for the probabilistically-delivered (risky) choice. This effect was blocked by co-administration of an oxytocin antagonist, confirming that actions on OT receptors in the brain mediated the attenuated risk taking (Tapp et al., 2020).

Choices between guaranteed vs. probabilistic rewards reflect one aspect of risk-taking behavior. Many real-world decisions, however, involve the possibility of explicit harm or loss, which are not well captured by the probabilistic discounting task. To examine potential contributions of OT to decision making under risk of explicit punishment, we used a risky decision-making task (RDT), in which rats choose between a small, “safe” food reward and a large, “risky” food reward that is accompanied by varying probabilities of mild footshock (Orsini et al., 2019; Simon et al., 2009). Previous work shows that preference for the large, risky reward in the RDT is associated with sensitivity to drugs of abuse and drug self-administration (Gabriel et al., 2019; Mitchell et al., 2014; Wheeler et al., 2023). Using this task, here we evaluated the effects of acute systemic administration of OT and an oxytocin antagonist (L-368,899 hydrochloride) on female and male rats’ behavior.

## Materials and Methods

### Subjects

Male and female Long-Evans rats (N=40; males=8, females=32; 55 days of age), were individually housed and allowed to acclimate to vivarium conditions for one week after arrival from Charles River Laboratories (Raleigh, NC). Rats were then food restricted to reach 85% of their free-feeding weight, followed by behavioral testing. During food restriction, rats’ target weights were adjusted upward by 5 g/week to account for growth. Rats were kept on a 12-hour light/dark cycle (lights on at 0700), maintained at a consistent temperature of 25°, fed on Teklad irradiated global 19% protein chow (2919), and had access to water *ad libitum*. Animal procedures were conducted in accordance with the University of Florida Institutional Animal Care and Use Committee and followed guidelines of the National Institutes of Health.

### Apparatus

Behavioral testing was conducted in eight standard operant chambers (Coulbourn Instruments). Chambers were contained in sound-attenuating cubicles, and were computer-controlled through Graphic State 4.0 software (Coulbourn Instruments). Locomotor activity was monitored via infrared motion detectors installed on the ceilings of the chambers. Each operant chamber was equipped with a food trough containing a photobeam sensor to detect nosepokes into the trough, two retractable levers (one on each side of the food trough), a feeder installed on the outside wall of the chamber and connected to the food trough to deliver 45 mg purified ingredient rodent food pellets (Test Diet; 1811155 5TUM), and a stainless steel floor grate connected to a shock generator that could deliver scrambled footshocks. Each sound-attenuating cubicle also included a house light mounted on the rear wall of the cubicle (outside of the operant chamber).

### Behavioral procedures

#### Risky decision-making task

Prior to testing in the RDT, rats were trained on a sequence of shaping protocols to learn how to retrieve food from the food trough, nosepoke in the food trough to initiate a trial, and press the levers to obtain food. Shaping procedures are described in detail in (Blaes et al., 2022). In the risky decision-making task (RDT; (Faraji et al., 2024; Orsini et al., 2019; Simon et al., 2009)), rats made discrete-trial choices between two levers. Each trial started with illumination of the house light and the food trough light. A nosepoke into the food trough during this time caused the food trough light to be extinguished and either one lever (on forced-choice trials) or both levers (on free-choice trials) to extend into the chamber. A failure to nosepoke within 10 s was counted as an omission. A press on one lever (the “small, safe” lever) yielded a single food pellet, whereas a press on the other lever (the “large, risky” lever) yielded two food pellets and was accompanied by varying probabilities of a mild footshock (1.0 s in duration). A failure to press either lever within 10 s resulted in termination of the trial and marked it as omission. Lever presses were followed by retraction of the lever(s), illumination of the food trough light, and delivery of food pellets. The food trough light was extinguished upon retrieval of the pellets or after 10 s, whichever came first. Trials were separated by an intertrial interval (ITI) in which the house light was extinguished. Each session was comprised of 5 blocks of trials, with different probabilities of shock accompanying the large reward in each block (0, 25, 50, 75, 100%). Figure 1 depicts a schematic of the RDT. Each block of trials began with 8 forced-choice trials (4 for each lever) in which only one lever was extended. The purpose of these trials was to remind rats of the probability of shock in that block. The forced-choice trials were followed by 10 free-choice trials in which both levers were extended. Sessions in the RDT were 60 minutes in duration and consisted of 90 trials, each 40 s long. The left/right positions of the small and large reward levers were randomized across rats, but remained consistent for each rat over the course of the task. Rats were trained on the RDT until stable choice performance emerged (see section 2.5, Data Analysis for definition of stable performance).

**Figure 1.**
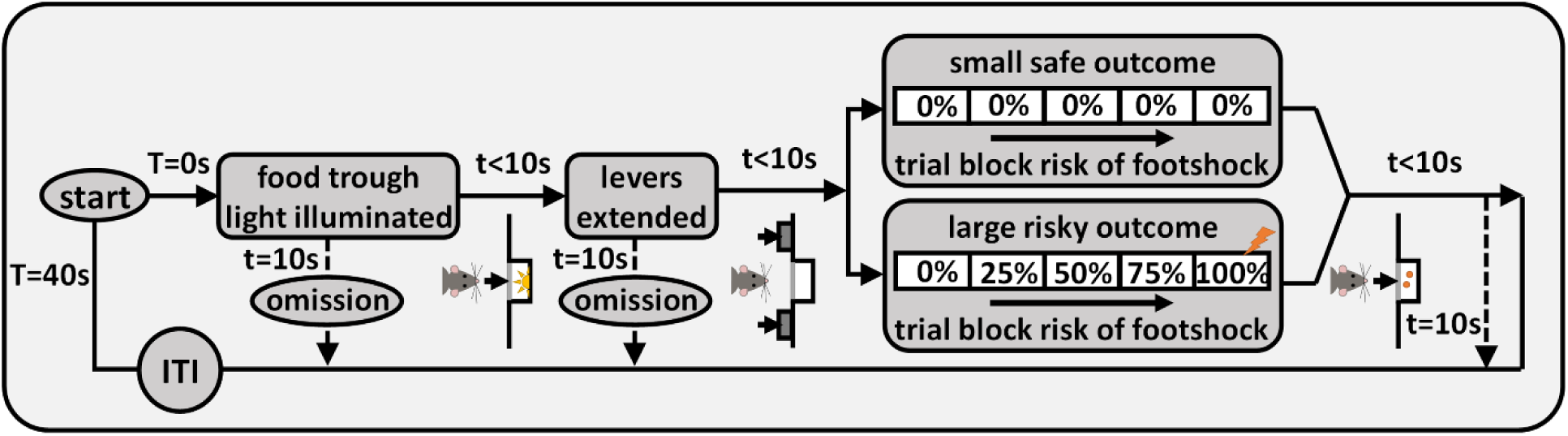
Risky decision-making task (RDT). Each trial was initiated by nosepoking into the food trough, and pressing a lever led to delivery of food pellet(s). If the large reward lever was chosen, delivery of this reward was accompanied by probabilistic shock delivery. There were five blocks of trials, and the probability of shock increased as blocks proceeded. “T” denotes the total time spent in the trial and “t” denotes the time spent in each state of the trial.

#### Drugs and drug administration

Both OT (Sigma-Aldrich, St. Louis, MO) and the OT receptor antagonist L-368,899 hydrochloride (MedChemExpress, Monmouth Junction, NJ) were dissolved in 0.9% physiological saline vehicle at concentrations of 3.0 mg/ml, and diluted to working solutions before use as needed. OT was freshly made on the day of experiment and L-368,899 was previously prepared and stored at −20 °C until use (L-368,899 stability at this storage condition was confirmed by manufacturer) on each injection day, when frozen aliquots were thawed at room temperature.

Rats received IP injections (1.0 ml/kg) of OT 15 min prior to testing, following a randomized, within-subject design (0, 0.3, 1.0, 3.0 mg/kg) such that each rat received each dose, with a 48 h washout period between successive doses. Upon completion of the OT administration schedule, rats continued training in the RDT until stable performance was again observed, at which point they proceeded to testing with L-368,899. Rats received IP injections (1.0 ml/kg) of L-368,899 40 minutes prior to testing, following a randomized, within-subject design (0, 1.0, 3.0 mg/kg) such that each rat received each dose, with a 48 h washout period between successive doses. The administration regimen, doses, and timing were chosen based on previous studies showing behavioral effects of both drugs following systemic administration (Carson, Hunt, et al., 2010; Hodges et al., 2019; Lefter et al., 2020; Smith et al., 2019; Tunstall et al., 2019). A total of n=16 female and n=8 male rats were tested with OT and L-368,899 in the RDT.

#### Shock reactivity threshold

To determine whether systemic administration of oxytocin or L-368,899 affects reactivity to shock, a separate cohort of rats (n=16, female) was tested in a shock reactivity threshold assay (Bonnet & Peterson, 1975; Orsini et al., 2017). Each rat was tested on the doses of OT and L-368,899 that elicited effects on risky decision making in the RDT. Shock reactivity threshold evaluations took place on four separate days, and under a pseudorandom design for IP injections (1.0 ml/kg) of saline, OT (0.3, 1.0 mg/kg), and L-368,899 (3.0 mg/kg), with at least a 48 h washout period between successive injections. For the test, rats were placed in a standard operant chamber and were initially administered a 400 *μ*A shock to attenuate spontaneous movement in the chamber. Five subsequent shocks (1.0 s in duration) were delivered at 10 s intervals, beginning at 50 *μ*A. Responses to the shocks were recorded by a trained observer. Shock reactivity criteria were defined as 1) flinch of a paw or a startle response, 2) elevation of one or two paws, 3) rapid movement of three or all paws. If out of the five shocks, fewer than three elicited any of the three reactivity behaviors, the intensity was increased by 25 *μ*A. This procedure continued until the rat was responsive to an intensity at least three out of five deliveries at that intensity, which was recorded as the shock reactivity threshold.

#### Reward magnitude discrimination task

To examine potential effects of oxytocin and L-368,899 on motivation to obtain the food reward or discriminate between small and large rewards in the RDT, the same rats tested for shock reactivity were tested on a reward magnitude discrimination task, using the doses of OT and L-368,899 that elicited effects on choice behavior in the RDT. This task was identical in structure to the RDT, except that the large reward was never accompanied by footshock. Rats were trained on the reward discrimination task until stable choice performance emerged (see section 2.5, Data Analysis for definition of stable performance). Reward discrimination was first evaluated with OT (0.3, 1.0 mg/kg and vehicle) on three separate days and under a pseudorandom design for IP injections (1.0 ml/kg) with at least a 48 h washout period between successive injections. Next, upon reemergence of stable behavior, reward discrimination was evaluated with L-368,899 (3.0 mg/kg and vehicle) on two separate days and under a pseudorandom design for IP injections (1.0 ml/kg) with at least a 48 h washout period between successive injections.

### Data analysis

Data were collected and processed using custom protocols and analysis templates in Graphic State 4.0. Statistical analyses were conducted and graphs created using GraphPad Prism 9. To evaluate stable performance, two-factor repeated measures analyses of variance (ANOVA, with session and trial block as within-subjects factors) were conducted on choice data across three consecutive sessions within each of the groups (male and female). Stability was defined as the absence of a significant main effect of session or session x trial block interaction.

The primary measure of interest in the RDT was the percentage of large, risky reward lever presses in each block (out of the total number of trials completed). For rats of each sex, choice behavior was evaluated using a two-factor repeated-measures ANOVA, with drug dose and trial block as within-subjects factors. In order to ensure sufficient parametric space in which to observe drug-induced changes in choice behavior, rats whose mean % choice of the large reward at the last 3 shock probabilities (50%, 75%, 100%) in vehicle sessions were less than 20% were eliminated in the corresponding OT (2 female rats) or L-368,899 (2 female rats) experiment. In the reward discrimination task, the primary measure of interest was the percentage of large reward lever presses in each block (out of the total number of trials completed). Choice behavior was evaluated using a two-factor repeated-measures ANOVA, with drug dose and trial block as within-subjects factors.

To examine the immediate effects of aversive outcomes on trial-by-trial choice behavior, win-stay/lose-shift analyses were conducted as in (Orsini et al., 2017). Trials on which choice of the large reward was not accompanied by a footshock were considered a “win”, and trials on which choice of the large reward was accompanied by a footshock were considered a “loss”. Win-stay was defined as when a win trial was followed by choice of the risky option on the next trial. Lose-shift was defined as when a lose trial was followed by a switch to the safe option on the next trial. Win-stay performance was calculated by dividing the number of win-stay trials by the total number of win trials, and lose-shift performance was calculated by dividing the number of lose-shift trials by the total number of lose trials. Both measures were calculated from free-choice trials only. Results were compared between drug doses using a one-factor repeated-measures ANOVA, with dose as the within-subjects factor. Shock reactivity threshold results were analyzed using a one-factor repeated-measure ANOVA, with drug condition (vehicle, OT, L-368,899) as the within-subjects factor.

Ancillary measures during the RDT such as locomotor activity, shock reactivity (locomotor activity during the shock delivery period), and omissions during free choice trials were analyzed using one-factor repeated-measure ANOVA, comparing data averaged across trial blocks. Latencies to press levers during forced-choice trials were used to assess motivation to pursue the outcomes associated with each choice. These data were analyzed using a three-factor repeated-measure ANOVA, with dose, lever identity, and trial block as within-subjects factors. For all analyses, the Greenhouse-Geisser correction was used to account for violations of sphericity in ANOVAs, and p values less than or equal to .05 were considered significant. Dunnett’s multiple comparison test was used for post-hoc analysis in the event ANOVA results were statistically significant.

### Transparency and openness

All procedures for statistical design, data analysis methods, and exclusion criteria are described in the manuscript. All data are available upon request from the corresponding author. This study’s design and analyses were not pre-registered.

## Results

### Risky decision-making task

#### Systemic administration of oxytocin

Rats were tested on the RDT until stable performance emerged (about 40 sessions). Next, rats received IP injections of OT using a randomized, within-subject design. In females, analysis of % choice of the large, risky reward using a two-factor repeated measures ANOVA (Dose x Shock Probability) revealed a main effect of Shock Probability (F(1.781, 21.377)=14.436, p<0.001), such that preference for the large reward declined as probability of shock increased. In addition, OT administration decreased the % choice of the large, risky reward (main effect of Dose, F(2.655, 31.860)=3.343, p=0.036), but there was no interaction between Dose and Shock Probability (F(4.851, 58.213)=1.520, p=0.199; Figure 2A). Follow-up two-factor ANOVAs comparing each dose with vehicle revealed a main effect of dose between the vehicle and 0.3 mg/kg (F(1, 12)=6.879, P=0.022), and the vehicle and 3 mg/kg (F(1, 12)=4.967, P=0.046) conditions, but no interaction between Dose and Shock Probability at either of these doses. In contrast to females, in males there was neither a main effect of Shock Probability (F(1.331, 9.317)=3.901, p=0.071) or Dose (F(2.376, 16.632)=1.914, p=0.174), nor was there an interaction between Dose and Shock Probability (F(2.571, 17.995)=0.984, p=0.412; Figure 2B).

**Figure 2.**
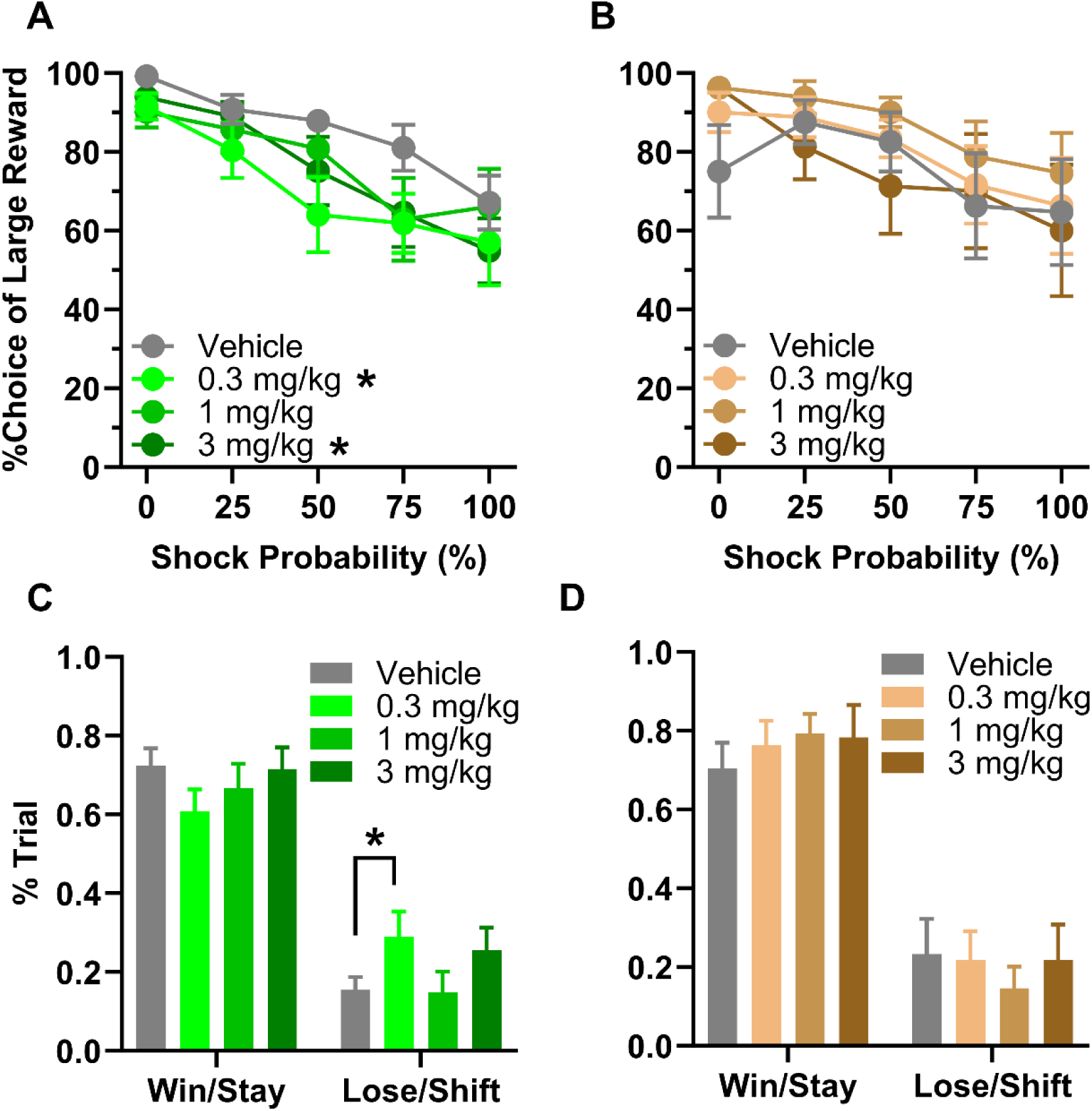
Systemic OT administration. **(A) Female choice behavior**. Females chose the large risky reward less frequently following administration of the 0.3 and 3.0 mg/kg doses of OT. **(B) Male choice behavior**. Males’ choice behavior was not affected by OT administration. **(C) Female win-stay/lose-shift behavior.** There was no effect of OT on win-stay performance, but the proportion of lose-shift trials was increased at the 0.3 mg/kg dose. **(D) Male win-stay/lose-shift behavior.** There was no effect of OT on either win-stay or lose-shift performance. Data are represented as means ± SEM.

Analysis of win-stay results using a one-factor repeated measures ANOVA (Dose) showed that the likelihood of making the same choice after a “win” was unaffected by OT in both females and males (see Table 1 for statistical results). In contrast, the same analysis conducted on lose-shift results showed that the likelihood of switching to the opposite choice after a loss (shock) was increased in females (F(2.605,31.262)=10.599, P<0.001; Figure 2C), but not in males (Figure 2D). Post-hoc Dunnett’s multiple comparisons tests revealed that the increase in lose-shift choices in females was significant at the 0.3 mg/kg dose (p=0.017), suggesting that OT increased rats’ sensitivity to the punished outcome.

**Table 1.** Choice strategies in the RDT.

***Win-Stay performance***
|  | OT |  | L-368,899 |  |
| --- | --- | --- | --- | --- |
|  | Female | Male | Female | Male |
| Dose | F(1.992,23.900)=1.597,<br>p=0.223 | F(1.671,11.696)=0.620,<br>p=0.527 | F(1.588,20.648)=1.106,<br>p=0.337 | F(1.632,11.426)=2.898,<br>p=0.103 |

***Lose-Shift performance***
|  | OT |  | L-368,899 |  |
| --- | --- | --- | --- | --- |
|  | Female | Male | Female | Male |
| Dose | F(2.605,31.262)=10.599,<br>P<0.001 | F(2.140,14.977)=0.814,<br>p=0.469 | F(1.727,20.724)=4.328,<br>p=0.031 | F(1.504,10.530)=0.462,<br>p=0.589 |

Latencies to press levers on forced-choice trials were analyzed using a three-factor repeated-measure ANOVA (Dose x Lever x Shock Probability). Latencies were longer at higher shock probabilities (Female: F(2.380,28.558)=19.124, p<0.001, Male: F(1.958,13.708)=6.874, P=0.009) and females (but not males) showed shorter latencies on the large compared to the small reward lever (Female: F(1, 12)=5.689, p=0.034). There were, however, no main effects or interactions involving Dose in either sex (Table 2).

**Table 2.**
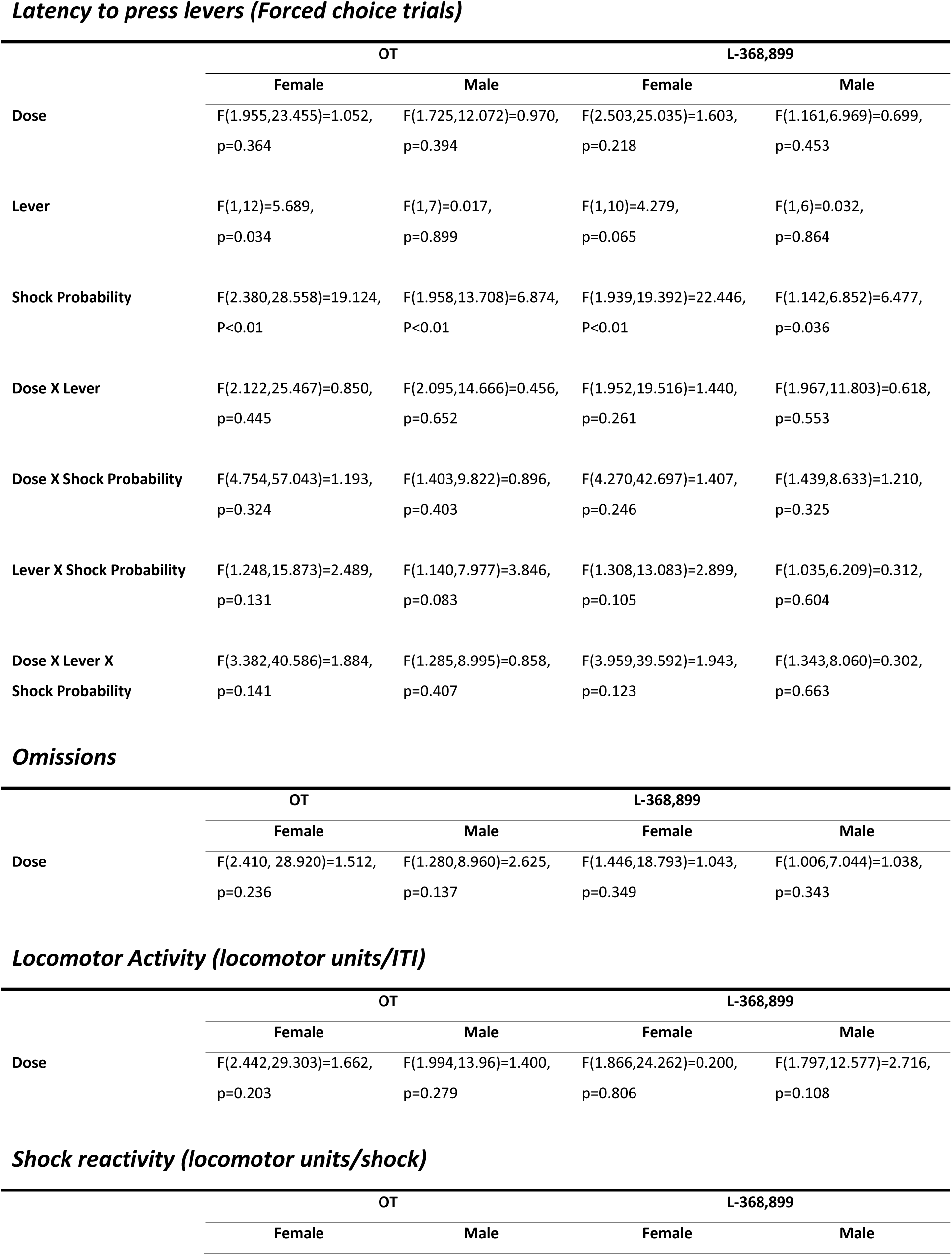
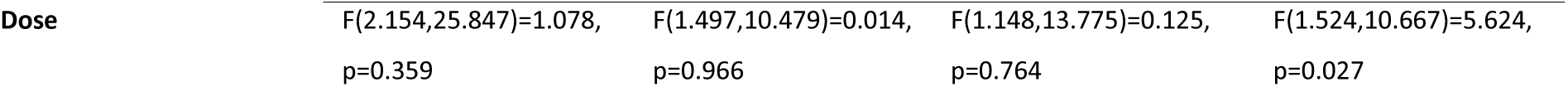
Ancillary measures in the RDT.

***Latency to press levers (Forced choice trials)***
|  | OT |  | L-368,899 |  |
| --- | --- | --- | --- | --- |
|  | Female | Male | Female | Male |
| **Dose** | F(1.955,23.455)=1.052,<br>p=0.364 | F(1.725,12.072)=0.970,<br>p=0.394 | F(2.503,25.035)=1.603,<br>p=0.218 | F(1.161,6.969)=0.699,<br>p=0.453 |
| **Lever** | F(1,12)=5.689,<br>p=0.034 | F(1,7)=0.017,<br>p=0.899 | F(1,10)=4.279,<br>p=0.065 | F(1,6)=0.032,<br>p=0.864 |
| **Shock Probability** | F(2.380,28.558)=19.124,<br>P<0.01 | F(1.958,13.708)=6.874,<br>P<0.01 | F(1.939,19.392)=22.446,<br>P<0.01 | F(1.142,6.852)=6.477,<br>p=0.036 |
| **Dose X Lever** | F(2.122,25.467)=0.850,<br>p=0.445 | F(2.095,14.666)=0.456,<br>p=0.652 | F(1.952,19.516)=1.440,<br>p=0.261 | F(1.967,11.803)=0.618,<br>p=0.553 |
| **Dose X Shock Probability** | F(4.754,57.043)=1.193,<br>p=0.324 | F(1.403,9.822)=0.896,<br>p=0.403 | F(4.270,42.697)=1.407,<br>p=0.246 | F(1.439,8.633)=1.210,<br>p=0.325 |
| **Lever X Shock Probability** | F(1.248,15.873)=2.489,<br>p=0.131 | F(1.140,7.977)=3.846,<br>p=0.083 | F(1.308,13.083)=2.899,<br>p=0.105 | F(1.035,6.209)=0.312,<br>p=0.604 |
| **Dose X Lever X Shock Probability** | F(3.382,40.586)=1.884,<br>p=0.141 | F(1.285,8.995)=0.858,<br>p=0.407 | F(3.959,39.592)=1.943,<br>p=0.123 | F(1.343,8.060)=0.302,<br>p=0.663 |

***Omissions***
|  | OT |  | L-368,899 |  |
| --- | --- | --- | --- | --- |
|  | Female | Male | Female | Male |
| **Dose** | F(2.410, 28.920)=1.512,<br>p=0.236 | F(1.280,8.960)=2.625,<br>p=0.137 | F(1.446,18.793)=1.043,<br>p=0.349 | F(1.006,7.044)=1.038,<br>p=0.343 |

***Locomotor Activity (locomotor units/ITI)***
|  | OT |  | L-368,899 |  |
| --- | --- | --- | --- | --- |
|  | Female | Male | Female | Male |
| **Dose** | F(2.442,29.303)=1.662,<br>p=0.203 | F(1.994,13.96)=1.400,<br>p=0.279 | F(1.866,24.262)=0.200,<br>p=0.806 | F(1.797,12.577)=2.716,<br>p=0.108 |

***Shock reactivity (locomotor units/shock)***
|  | OT |  | L-368,899 |  |
| --- | --- | --- | --- | --- |
|  | Female | Male | Female | Male |
| **Dose** | F(2.154,25.847)=1.078,<br>p=0.359 | F(1.497,10.479)=0.014,<br>p=0.966 | F(1.148,13.775)=0.125,<br>p=0.764 | F(1.524,10.667)=5.624,<br>p=0.027 |

A one-factor repeated measure ANOVA (Dose) was used to evaluate locomotor activity during the RDT. OT administration did not affect locomotor activity during the ITI, or during shock deliveries in the task (locomotor activity during shock delivery) in either females or males (Table 2). Analysis of the number of free-choice trial omissions also revealed no effects of Dose (Table 2).

#### Systemic administration of L-368,899

After completion of the OT administration regimen, rats continued testing on the RDT until stable performance emerged again. Rats then received IP injections of L-368,899 hydrochloride using a randomized within-subject design. In females, analysis of % choice of the large, risky reward using a two-factor, repeated measures ANOVA (Dose x Shock Probability) revealed a main effect of Shock Probability (F(2.601, 33.807)=18.220, p<0.001), such that preference for the large reward declined as probability of shock increased. Similar to the OT administration results, systemic administration of L-368,899 decreased % choice of the large, risky reward (main effect of Dose, F(1.647, 21.416)=4.304, p=0.033), but there was no interaction between Dose and Shock Probability (F(3.470, 43.806)=1.046, p=0.389; Figure 3A). Post-hoc comparisons revealed a main effect of dose between the vehicle and 3.0 mg/kg (F(1, 13)=11.130, P=0.005) conditions, but no interaction between Dose and Shock Probability. In contrast to females, in males there were no significant effects of Shock Probability (F(2.089, 14.624)=3.425, p=0.059) or Dose (F(1.481, 10.369)=0.493, p=0.570), nor was there an interaction between the two variables (F(2.742, 19.193)=1.533, p=0.239; Figure 3B).

**Figure 3.**
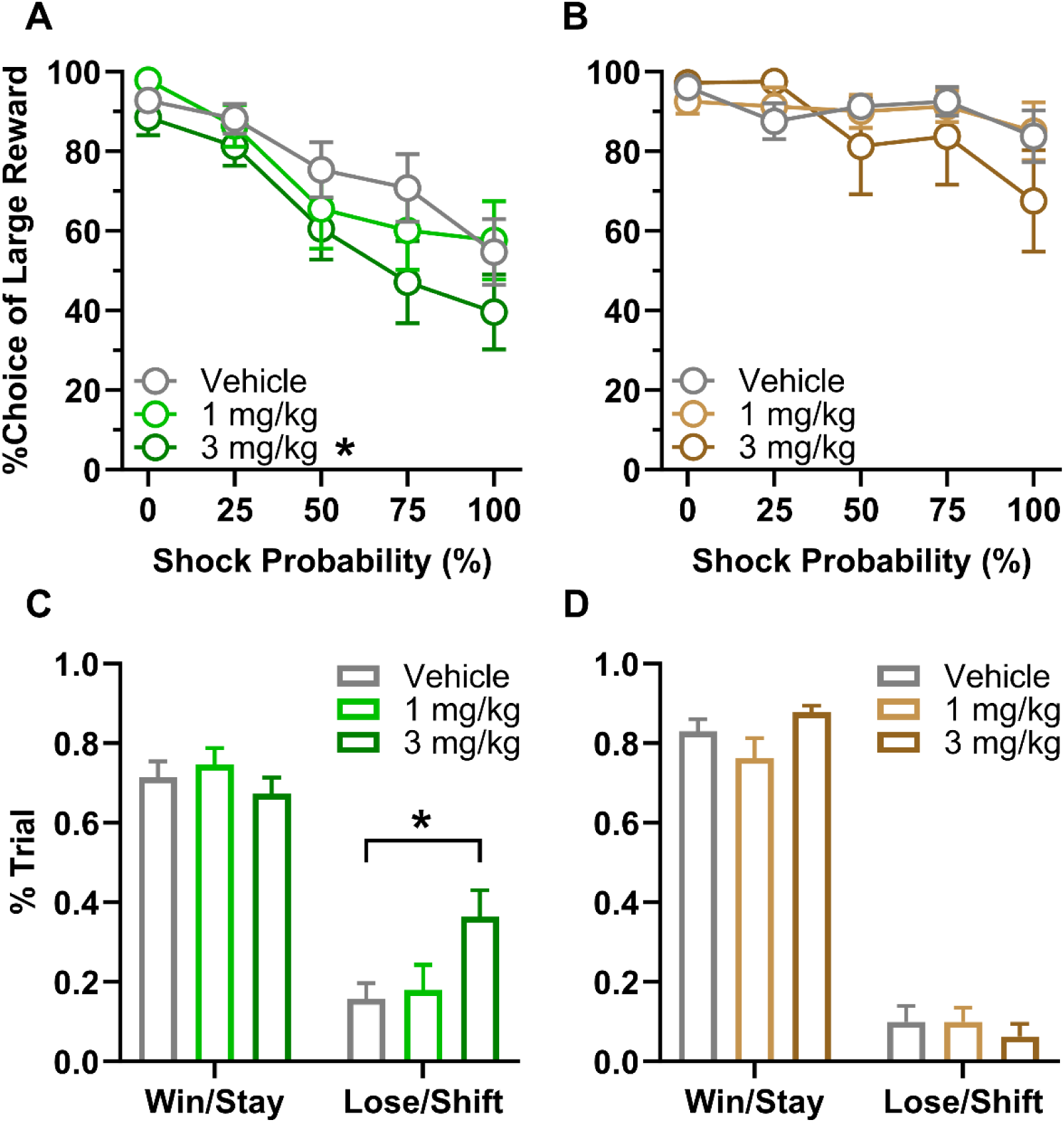
Systemic L-368,899 administration. **(A) Female choice behavior**. Females chose the large risky reward less frequently following administration of the 3.0 mg/kg dose of L-368,899. **(B) Male choice behavior**. Males’ choice behavior was not affected by L-368,899 administration. **(C) Female win-stay/lose-shift behavior.** There was no effect of L-368,899 on win-stay performance, but the proportion of lose-shift trials was increased at the 3.0 mg/kg dose. **(D) Male win-stay/lose-shift.** There was no effect of L-368,899 on either win-stay or lose-shift performance. Data are represented as means ± SEM.

Analysis of win-stay results using a one-factor repeated measures ANOVA (Dose) showed that the likelihood of making the same choice after a “win” was unaffected by Dose in both females and males (Table 1). In contrast, the same analysis conducted on lose-shift results showed that the likelihood of switching to the opposite choice after a loss (shock) was increased as the dose increased in females (F(1.727,20.724)=4.328, p=0.031; Figure 3C), but not in males (Figure 3D). Post-hoc analysis revealed that the increase in lose-shift choices in females was significant at the 3.0 mg/kg L-368,899 dose (p=0.03).

Latencies to press levers on forced-choice trials were analyzed using a three-factor repeated-measure ANOVA (Dose x Lever x Shock Probability). Latencies were longer at higher shock probabilities (Female: F(1.939,19.392)=22.446, P<0.001, Male: F(1.142,6.852)=6.477, p=0.036). There were, however, no other significant main effects or interactions (Table 2).

A one-factor repeated measure ANOVA (Dose) was used to evaluate locomotor activity during the RDT. L-368,899 administration did not affect locomotor activity during the ITI in either females or males (Table 2). The same analysis on shock reactivity during the task (locomotor activity during shock delivery) revealed a main effect of Dose in males, such that activity during shock delivery was reduced at the highest dose (F(1.524,10.667)=5.624, p=0.027), but there was no effect of Dose in females (Table 2). Analysis of the number of free-choice trial omissions revealed no main effect of Dose in either females or males (Table 2).

### Shock reactivity threshold

As both OT and L-368,899 reduced female rats’ choice of the large, risky reward, we tested a separate cohort of females in a more sensitive assay of shock reactivity using the doses of OT (0.3 and 3.0 mg/kg) and L-368,899 (3.0 mg/kg) that elicited effects on choice behavior in the RDT, to determine whether the effects of the drugs could be due to increases in shock sensitivity (Figure 4). Comparison of reactivity thresholds using a one-way ANOVA (Drug Condition) revealed a main effect of Drug Condition (F(2.007, 30.100)=3.824, p=0.033), with post-hoc tests showing a significant increase in shock reactivity threshold following both OT (3.0 mg/kg, p=0.045) and L-368,899 (3.0 mg/kg, p=0.033) compared to vehicle, but no effects at 0.3 mg/kg OT (p=0.675).

**Figure 4.**
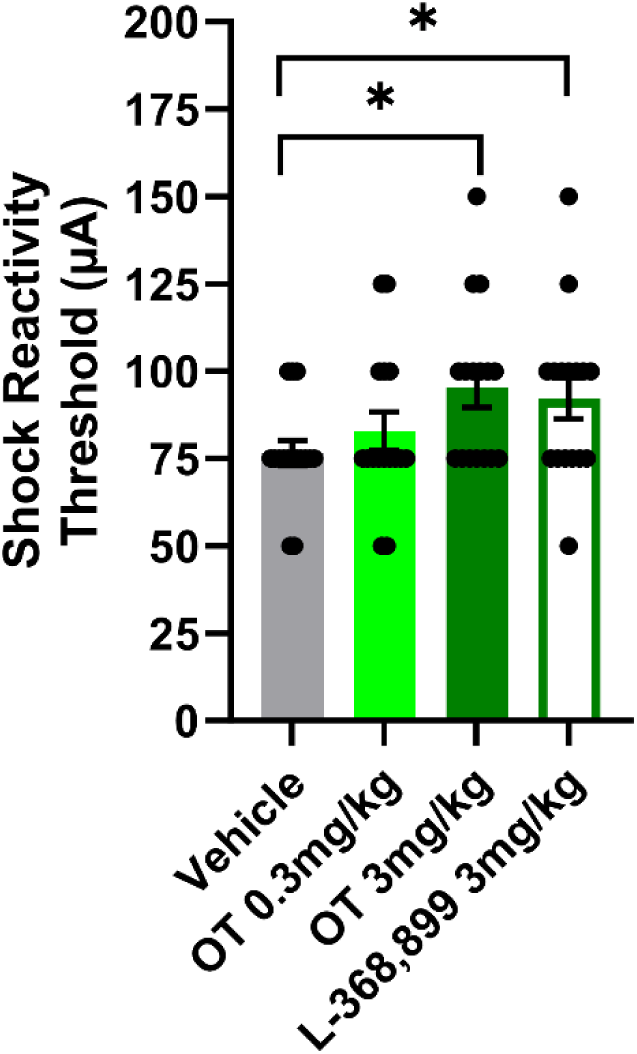
Shock reactivity threshold. Both OT (3.0 mg/kg) and L-368,899 (3.0 mg/kg) increased shock reactivity thresholds in females. Bars represent means + SEM, and points represent individual data from each rat.

### Reward magnitude discrimination

#### Systemic administration of oxytocin

The same cohort of female rats used to evaluate shock reactivity was then tested on the reward magnitude discrimination task until stable performance emerged (6 sessions). Next, rats received IP injections of OT using a randomized within-subject design. Analysis of % choice of the large reward using a two-factor repeated measures ANOVA (Dose x Trial Block) revealed no main effect of Dose (F(1.346, 150.78)=1.086, p=0.319, Figure 5A). In addition, the choice preference of female rats remained consistent during the session such that preference for the large reward did not change as the trials proceeded (Trial Block, F(2.540, 142.23)=1.282, p=0.283, Dose x Trial Block, F(3.599, 100.78)=1.101, p=0.358). Analysis of the number of free-choice trial omissions also showed that task participation was not affected by OT (Table 3). These results confirm that reduced choice of large reward in RDT following systemic administration of OT was not associated with altered motivation for food.

**Figure 5.**
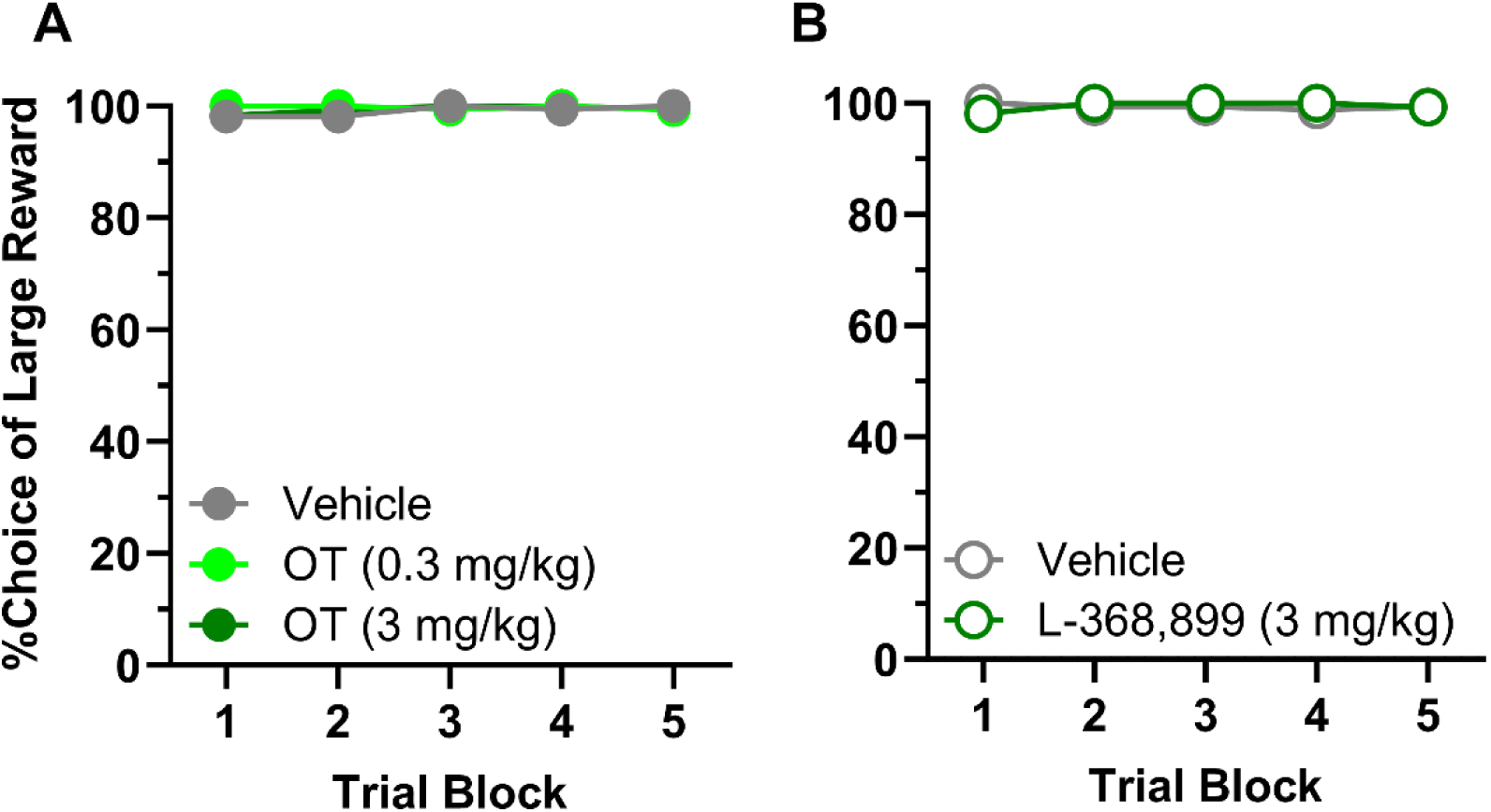
Reward magnitude discrimination. **(A) Systemic OT administration.** There were no effects on females’ choice of the large reward following administration of the 0.3 and 3.0 mg/kg doses of OT. **(B) Systemic L-368,899 administration.** There were no effects on females’ choice of large reward following administration of the 3.0 mg/kg dose of L-368,899. Data are represented as means ± SEM.

**Table 3.**
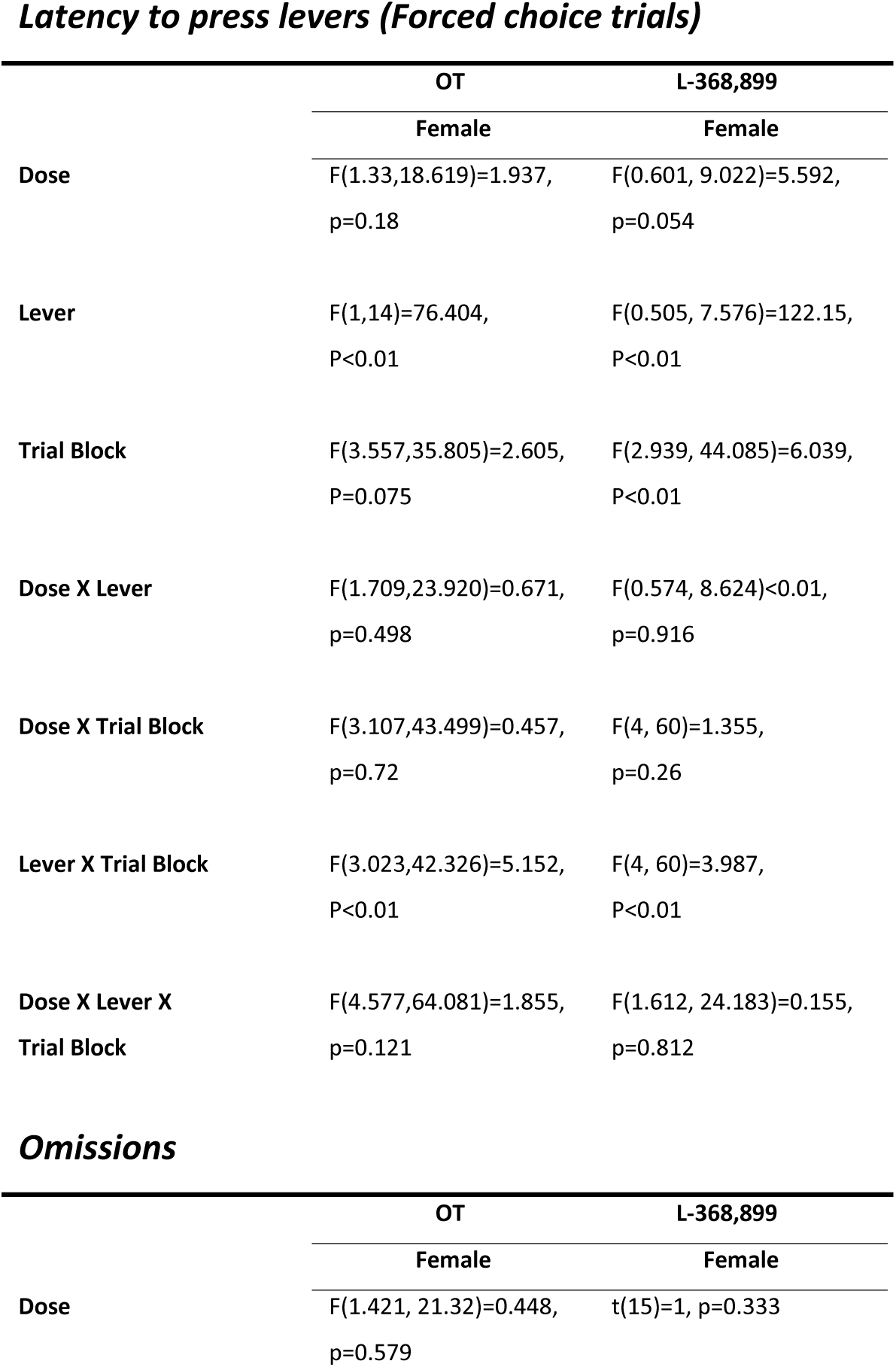
Ancillary measures in the reward discrimination task.

***Latency to press levers (Forced choice trials)***
|  | OT | L-368,899 |
| --- | --- | --- |
|  | Female | Female |
| **Dose** | F(1.33,18.619)=1.937,<br>p=0.18 | F(0.601, 9.022)=5.592,<br>p=0.054 |
| **Lever** | F(1,14)=76.404,<br>P<0.01 | F(0.505, 7.576)=122.15,<br>P<0.01 |
| **Trial Block** | F(3.557,35.805)=2.605,<br>P=0.075 | F(2.939, 44.085)=6.039,<br>P<0.01 |
| **Dose X Lever** | F(1.709,23.920)=0.671,<br>p=0.498 | F(0.574, 8.624)<0.01,<br>p=0.916 |
| **Dose X Trial Block** | F(3.107,43.499)=0.457,<br>p=0.72 | F(4, 60)=1.355,<br>p=0.26 |
| **Lever X Trial Block** | F(3.023,42.326)=5.152,<br>P<0.01 | F(4, 60)=3.987,<br>P<0.01 |
| **Dose X Lever X**<br>**Trial Block** | F(4.577,64.081)=1.855,<br>p=0.121 | F(1.612, 24.183)=0.155,<br>p=0.812 |

***Omissions***
|  | OT | L-368,899 |
| --- | --- | --- |
|  | Female | Female |
| **Dose** | F(1.421, 21.32)=0.448,<br>p=0.579 | t(15)=1, p=0.333 |

Latencies to press levers on forced-choice trials in this task were analyzed using a three-factor repeated-measure ANOVA (Dose x Lever x Trial Block). Latencies were longer when pressing the small reward lever (F(1,14)=76.404, P<0.001). There was an interaction between the Lever and Trial Block such that the latency to press the small reward lever increased as the trial blocks proceeded (F(3.023,42.326)=5.152, P<0.001). There were, however, no other significant main effects or interactions concerning Dose (Table 3).

#### Systemic administration of L-368,899

After completion of the OT administration regimen, rats continued testing on the reward magnitude discrimination task until stable performance emerged again. Rats then received IP injections of L-368,899 hydrochloride using a randomized within-subject design. Analysis of % choice of the large reward using a two-factor repeated measures ANOVA (Dose x Trial Block) revealed no main effect of Dose (F(1, 15)=0.081, p=0.78, Figure 5B). In addition, the choice preference of female rats remained consistent during the session such that preference for the large reward did not change as the trials proceeded (Trial Block, F(2.169, 32.541)=0.477, p=0.640, Dose x Trial Block, F(2.436, 36.538)=2.156, p=0.121). Analysis of the number of free-choice trial omissions also revealed that task participation was not affected by L-368,899 (Table 3). These results confirm that reduced choice of large reward in RDT following systemic administration of L-368,899 was not associated with altered motivation for food.

Latencies to press levers on forced-choice trials were analyzed using a three-factor repeated-measure ANOVA (Dose x Lever x Trial Block). On forced-choice trials, latencies were longer when pressing the small reward lever (Lever, F(0.505, 7.576)=122.15, P<0.01) and in later trials during a session (Trial Block, F(2.939, 44.085)=6.039, P<0.01). There was also an interaction between the Lever and Trial Block such that the latency to press the small reward lever increased as the trial blocks proceeded (F(4, 60)=3.987, P<0.01). There were, however, no other significant main effects or interactions concerning Dose (Table 3).

## Discussion

Despite well-documented roles in reward- and stress-related behaviors, the role of OT signaling in cost-benefit decision making and particularly risky decision making is less well understood. Here we show that OT signaling modulates decision making under risk of explicit punishment, and that this effect is sex dependent. In females (but not males), both systemic OT and systemic blockade of OT receptors via the selective antagonist L-368,899 reduced preference for large food rewards accompanied by risk of footshock punishment over small, unpunished rewards (i.e., reduced risk taking). These effects were not due to elevated sensitivity to the footshock punishment, as both drugs actually reduced shock reactivity at doses that attenuated risk taking. Moreover, the reduced preference for the large reward was not due to reduced motivation for food or impaired ability to discriminate between the small and large reward, as the results of the reward magnitude discrimination task revealed that neither OT nor its antagonist affected preference for the large reward or the number of omitted trials. Given that maladaptive risk taking accompanies (and in some cases is a central feature of) numerous neuropsychiatric conditions, these results suggest that targeting OT signaling could be a therapeutic strategy for addressing decision making deficits in such conditions.

### Decreased risk taking with systemic OT

As a neurohormone with actions throughout the periphery and the brain, OT has effects on a number of behavioral processes that could contribute to choice behavior in the RDT. For example, OT can modulate several measures of pain sensitivity. Research in humans and rodents shows that endogenous OT inhibits both somatic and visceral nociception (Goodin et al., 2015). Exogenously, intrathecal and intranasal OT have proven efficacy in chronic and acute pain management in humans (Lussier et al., 2019). Administration of OT to women during labor is a common practice (Uvnäs-Moberg, 2023), intranasal and intravenous OT is effective in attenuating back pain and migraines (Mekhael et al., 2023), and patients with irritable bowel syndrome show higher threshold for colonic visceral pain with OT treatments (Louvel et al., 1996). In rodents, peripherally administered OT reduces pain sensitivity (de Araujo et al., 2014; Schorscher-Petcu et al., 2010). Further studies show that the analgesic actions of OT are modulated at spinal, supraspinal, and peripheral levels (Xin et al., 2017), and that both neuronal and humoral pathways are active in OT modulation of pain (Nishimura et al., 2022). The finding in the current study that OT increased the intensity at which rats reacted to the shock during the shock reactivity test is consistent with this previous work. Importantly, however, this reduction in shock sensitivity does not account for the OT-induced reduction in preference for the large, risky reward in the RDT, suggesting that the effects of OT on risk taking are separate from (and even in spite of) its analgesic effects.

Another possible mechanism for the decrease in risk taking following systemic OT concerns effects on food motivation and/or valuation. Exogenous OT can reduce food intake in animals and humans (Lawson, 2017). OT exerts its anorexigenic effects via diverting resources to competing motivations such as reproduction, fear, or stress, and via interactions with satiation and reward signaling, nutrient-specific effects on food intake, and social factors (Liu et al., 2021). Multiple brain regions facilitate OT modulation of food intake and appetitive behavior. In rats, direct OT infusion into the basolateral and central nuclei of the amygdala reduced deprivation-induced food intake (Klockars et al., 2018), and intracerebroventricular administration of OT reduced phasic dopaminergic neuron activity in the VTA in response to sucrose-predictive cues (Liu et al., 2020). In humans, fMRI studies showed that intranasal administration of OT enhanced the activity of brain regions responsible for cognitive control (e.g., frontal cortex) and subsequently reduced food intake (Spetter et al., 2018). Such OT-induced reductions in food motivation could in theory account for the reduction in rats’ preference for the large, risky reward in the RDT, e.g., by reducing its value relative to the small, safe reward; however, several lines of evidence argue against this interpretation. First, several prior studies show that reducing food motivation via either acute or 24 h pre-feeding does not affect rats’ risk taking behavior in the RDT (though it does cause an increase in the number of omitted trials), suggesting that relatively acute changes in hunger and satiation do not strongly drive choice preference in this task (Orsini et al., 2016; Simon et al., 2009). Second, the reduction in choice of the large reward induced by OT was most robust in the presence of shock, and was minimal or absent in the first block of trials, when no shock was present. Third, OT produced an increase in the proportion of lose/shift trials, but had no effects on win/stay trials, suggesting that its effects were primarily on aversive rather than appetitive motivating factors in the task. Finally, we tested the rats on a reward discrimination task which is identical to RDT in design except that no punishment accompanied the large reward. The results revealed no effect of OT on choice behavior in this task, nor was there an increase in trial omissions, strongly suggesting that the decrease in choice of large reward in the RDT was not due to anorexic effects of OT. Notably, these findings are consistent with those of Tapp et al., who found that systemic OT administration (at 6.0 mg/kg, which is twice the highest dose used here) failed to affect choice behavior in a probability discounting task that was similar in design to the RDT but in which the large reward was associated with variable probabilities of omission (Tapp et al., 2020). The absence of effects of OT on choices between small and large reward when the “cost” associated with the large reward was its possible omission suggests that OT’s actions on choice behavior in the RDT were due to the specific nature of the cost (i.e., punishment). The fact that OT selectively affected lose/shift and not win/stay trials is consistent with this interpretation.

OT actions across multiple brain regions can influence a variety of aversively-motivated behaviors, including contextual fear, cued fear, socially transmitted fear, and fear discrimination (Olivera-Pasilio & Dabrowska, 2020). The amygdala in particular is the most frequently reported brain region in the context of OT-induced modulation of aversive states in both human and animal studies. In humans, intranasal OT has proven effective in accelerating fear extinction in patients with posttraumatic stress disorder, generalized anxiety, and phobias, presumably, by both decreasing amygdala activity during negative-valenced processes and increasing amygdala activity promoting approach behavior during social and appetitive processes (Baldi et al., 2021; Wang et al., 2017). In rodents, both intranasal OT and direct infusion of OT into the central amygdala facilitated fear extinction (Baldi et al., 2021). Interestingly, despite these overall anxiolytic effects, OT acting in the central nucleus of the amygdala can facilitate a switch from passive to active coping mechanisms in the presence of an escapable threat (i.e., OT can promote avoidance behavior) (Terburg et al., 2018). Such facilitation of avoidance could account for the reduction in choice of the large, risky reward induced by OT.

Systemic administration of OT can facilitate dopamine release in the nucleus accumbens (NAc) via activating dopaminergic neurons in ventral tegmental area that project to NAc (Baracz & Cornish, 2016; Kohli et al., 2019). Dopamine signaling – particularly in the NAc – strongly modulates choice performance in the RDT. Risk aversion is associated with higher levels of D2 dopamine receptor mRNA in the NAc, and both systemic and intra-NAc activation of D2 receptors reduces risk taking (Blaes et al., 2018; Mitchell et al., 2014; Simon et al., 2011). In addition to facilitating dopamine release in NAc, systemic OT may modulate risk taking via enhancing activation of D2 dopamine and OT receptor heteromers in the NAc shown previously to be involved in pair bonding behavior (Fuxe et al., 2012). As such, it is possible that OT-induced increases in NAc dopamine and OT signaling could account at least in part for its actions in the RDT.

### Decreased risk taking with systemic L-368,899

Given that systemic administration of OT reduced rats’ preference for the large, risky reward, it was somewhat surprising that systemic administration of the selective OT receptor antagonist L-368,899 had the same effect (again, in females but not in males). At face value these two sets of data could appear contradictory; however, it is important to note that whereas OT acts at arginine vasopressin (AVP) receptors in addition to OT receptors, L-368,899 is relatively selective (>40-fold) for OT receptors. One mechanism that might explain L-368,899 effects is the well documented OT-AVP cross talk (Qiu et al., 2014; Rae et al., 2022; Schorscher-Petcu et al., 2010; Song & Albers, 2018). For example, AVP receptor-expressing neurons in the medial portion of the central nucleus of the amygdala (CeM) are activated by BLA excitatory inputs and are critical in processing of aversive stimuli by gating inputs from the lateral portion of the central nucleus of the amygdala (CeL), which in turn are modulated by OT receptors (Huber et al., 2005). At least in male rats, the BLA modulates encoding of punishment in the RDT, as lesions of this structure increase risk taking (Orsini et al., 2015) and optogenetic inhibition of BLA selectively during receipt of the large, punished large outcome has the same effect (Orsini et al., 2017). If BLA projections to CeM were involved in biasing behavior away from risky options, then OT receptor blockade could disinhibit the AVP-expressing neurons in CeM (which are normally inhibited by OT-expressing neurons projecting from CeL) that are activated by BLA input, facilitating these BLA-derived signals and ultimately decreasing risk taking. This of course raises the question as to why systemic OT did not cause the opposite effect by enhancing the inhibitory influence of OT receptor-expressing neurons in CeL onto AVP receptor-expressing neurons in CeM; however, OT at high concentrations can activate AVP receptors as well, and thus it is likely that exogenous OT, through non-selective binding, can act on both CeL and CeM neurons, resulting in increased net output from CeM. Future work is needed to address the specific roles of AVP and AVP receptor signaling in risky decision making.

It is important to note that the effects of L-368,899 on aspects of RDT performance aside from choice preference were similar to those of OT, in that it increased the proportion of lose/shift trials and had no effect on choice behavior in the absence of shock. In addition, like OT, it increased the threshold for shock reactivity, suggesting that its effects on preference for the large, risky reward were not due to enhanced shock sensitivity. Moreover, the fact that the effect of L-368,899 in the shock reactivity assay was in the same direction as that produced by OT suggests that it acted via a similar mechanism across both tasks.

### Sex differences

Unlike female rats, males did not show any change in choice behavior in the RDT following systemic administration of either OT or L-368,899. It is possible that sex differences in OT and AVP signaling could account for the absence of OT and OT antagonist effects on risky decision making in male rats. Indeed, there is evidence for sex differences in the contributions of OT and AVP systems to some forms of social decision making (Dumais & Veenema, 2016; Jiang & Platt, 2018; Lu et al., 2019; Smith et al., 2017), and it is possible that this sexual dimorphism extends to non-social decision-making as well. Another possibility is that male rats are less sensitive to OT and L-368,899 at the doses and shock intensities used in the present study. Specifically, OT and OT receptor antagonism increased lose-shift trials in females, but had no effect on lose-shift performance in males, indicating a lack of change in aversion toward the probabilistic shock in males. If this were the case, it is likely that higher shock intensities or higher doses of OT and L-368,899 would elicit changes in the RDT choice behavior in males. Future work is needed to address the role of parameter sensitivity in the effects of OT signaling modulation on risky decision making in males.

### Future directions

The similar pattern of reduced risk-taking behavior resulting from systemic administration of OT and L-368,899 speaks to the complexity of OT signaling. Exogenous OT has proven effective in reducing anxiety and drug-seeking behavior (Baracz et al., 2022; King et al., 2020; Leong et al., 2018; Neumann & Slattery, 2016; Rhianne et al., 2023; Yoshida et al., 2009), but there are numerous unanswered questions concerning the mechanisms of these effects. One of the important directions necessary to elevate the efficacy of OT-related treatments is to investigate cross talk between OT and AVP systems, which plays a crucial role in emotional regulation including fear responses (Gozzi et al., 2017; Stevens, 2005; Viviani & Stoop, 2008). Investigating the role of AVP receptor signaling in facilitating/gating exogenous OT and OT antagonist effects could be instrumental to identification of novel therapeutic targets. Another possible translationally-relevant direction would be to evaluate the consequences of chronic administration. OT influences the HPA axis, and chronic OT administration is reported to elicit adverse behavioral effects associated with increased ACTH and corticosterone levels (Jurek & Meyer, 2020). The effects of chronic OT are dose dependent, as low-dose chronic OT in mice decreased anxiety-like behavior, whereas high-dose chronic OT had the opposite effects (Peters et al., 2014). Deciphering the dynamics of oxytocinergic signaling in relation to other hormonal systems appear to be a necessary step in developing OT-based therapeutics.

While for therapeutic purposes, OT, AVP, or their antagonists are administered peripherally, a large body of research has investigated OT and AVP signaling via intracerebral routes of administration. A number of studies show differential behavioral effects of OT based on the route of administration (Kou et al., 2021; Tapp et al., 2020; Valstad et al., 2016; Yao & Kendrick, 2022), which underscores the importance of considering this factor as a consequential variable in both experimental design and translational utility.

## Summary and Conclusions

Here we showed that systemic administration of both OT and the OT receptor antagonist L-368,899 reduced risk taking in female but not male rats. Control experiments showed that these reductions in risk taking were due neither to increased sensitivity to the footshock punishment, nor to reduced attention to or motivation to obtain the food rewards. To our knowledge, these results are the first to investigate the role of OT in decision making under risk of explicit punishment. Given that maladaptive increases in risky behavior accompany numerous psychiatric disorders, these results suggest that further research on the role of OT signaling in risky decision making could ultimately yield therapeutic benefits.

## References

Albin-Brooks, C., Nealer, C., Sabihi, S., Haim, A., & Leuner, B. (2017). The influence of offspring, parity, and oxytocin on cognitive flexibility during the postpartum period. Horm Behav, 89, 130–136. 10.1016/j.yhbeh.2016.12.015

Baldi, E., Costa, A., Rani, B., Passani, M. B., Blandina, P., Romano, A., & Provensi, G. (2021). Oxytocin and Fear Memory Extinction: Possible Implications for the Therapy of Fear Disorders? Int J Mol Sci, 22(18). 10.3390/ijms221810000

Ballas, H. S., Wilfur, S. M., Freker, N. A., & Leong, K.-C. (2021). Oxytocin attenuates the stress-induced reinstatement of a lcohol-seeking in male rats: Role of the central amygdala Biomedicines, 9(12), 1919–1928. 10.3390/biomedicines9121919

Baracz, S. J., & Cornish, J. L. (2016). The neurocircuitry involved in oxytocin modulation of methamphetamine addiction. Frontiers in Neuroendocrinology, 43, 1–18. 10.1016/j.yfrne.2016.08.001

Baracz, S. J., Everett, N. A., McGregor, I. S., & Cornish, J. L. (2014). Oxytocin in the nucleus accumbens core reduces reinstatement of methamphetamine-seeking behaviour in rats. Addiction Biology, 21(2), 316–325. 10.1111/adb.12198

Baracz, S. J., Robinson, K. J., Wright, A. L., Turner, A. J., McGregor, I. S., Cornish, J. L., & Everett, N. A. (2022). Oxytocin as an adolescent treatment for methamphetamine addiction after early life stress in male and female rats. Neuropsychopharmacology, 47(8), 1561–1573. 10.1038/s41386-022-01336-y

Baracz, S. J., Rourke, P. I., Pardey, M. C., Hunt, G. E., & McGregor, I. S. (2012). Oxytocin directly administered into the nucleus accumbens core or subthalamic nucleus attenuates methamphetamine-induced conditioned place preference. Behavioral Brain Research, 228(1), 185–193. 10.1016/j.bbr.2011.11.038

Bartz, J. A., Zaki, J., Bolger, N., Hollander, E., Ludwig, N. N., Kolevzon, A., & Ochsner, K. N. (2010). Oxytocin selectively improves empathic accuracy. Psychol Sci, 21(10), 1426–1428. 10.1177/0956797610383439

Baumgartner, T., Heinrichs, M., Vonlanthen, A., Fischbacher, U., & Fehr, E. (2008). Oxytocin shapes the neural circuitry of trust and trust adaptation in humans. Neuron, 58(4), 639–650. 10.1016/j.neuron.2008.04.009

Blaes, S. L., Orsini, C. A., Mitchell, M. R., Spurrell, M. S., Betzhold, S. M., Vera, K., Setlow, B. (2018). Monoaminergic modulation of decision-making under risk of punishment in a rat model. Behav Pharmacol, 29(8), 745–761. 10.1097/fbp.0000000000000448

Blaes, S. L., Shimp, K. G., Betzhold, S. M., Setlow, B., & Orsini, C. A. (2022). Chronic cocaine causes age-dependent increases in risky choice in both males and females. Behavioral Neuroscience, 136, 243–263. 10.1037/bne0000509

Bonnet, K. A., & Peterson, K. E. (1975). A modification of the jump-flinch technique for measuring pain sensitivity in rats. Pharmacol Biochem Behav, 3(1), 47–55. 10.1016/0091-3057(75)90079-9

Bozorgmehr, A., Alizadeh, F., Sadeghi, B., Shahbazi, A., Norouzi Ofogh, S., Joghataei, M. T., … Ghadirivasfi, M. (2019). Oxytocin moderates risky decision-making during the Iowa Gambling Task: A new insight based on the role of oxytocin receptor gene polymorphisms and interventional cognitive study. Neuroscience Letters, 708, 134328. 10.1016/j.neulet.2019.134328

Carcea, I., Caraballo, N. L., Marlin, B. J., Ooyama, R., Riceberg, J. S., Mendoza Navarro, J. M., … Froemke, R. C. (2021). Oxytocin neurons enable social transmission of maternal behaviour. Nature, 596(7873), 553–557. 10.1038/s41586-021-03814-7

Carson, D. S., Cornish, J. L., Guastella, A. J., Hunt, G. E., & McGregor, I. S. (2010). Oxytocin decreases methamphetamine self-administration, methamphetamine hyperactivity, and relapse to methamphetamine-seeking behaviour in rats. Neuropharmacology, 58, 38–43. 10.1016/j.neuropharm.2009.06.018

Carson, D. S., Hunt, G. E., Guastella, A. J., Barber, L., Cornish, J. L., Arnold, J. C., McGregor, I. S. (2010). Systemically administered oxytocin decreases methamphetamine activation of the subthalamic nucleus and accumbens core and stimulates oxytocinergic neurons in the hypothalamus. Addiction Biology, 15, 448–463. 10.1111/j.1369-1600.2010.00247.x

Chen, S., Yang, P., Chen, T., Su, H., Jiang, H., & Zhao, M. (2020). Risky decision-making in individuals with substance use disorder: A meta-analysis and meta-regression review. Psychopharmacology, 237, 1893–1908. 10.1007/s00213-020-05506-y

Cox, B. M., Young, A. B., See, R. E., & Reichel, C. M. (2013). Sex differences in methamphetamine seeking in rats: Impact of oxytocin. Psychoneuroendocrinology, 38(10), 2343–2353. 10.1016/j.psyneuen.2013.05.005

de Araujo, A. D., Mobli, M., Castro, J., Harrington, A. M., Vetter, I., Dekan, Z., Alewood, P. F. (2014). Selenoether oxytocin analogues have analgesic properties in a mouse model of chronic abdominal pain. Nature Communications, 5(1), 3165. 10.1038/ncomms4165

De Dreu, C. K. (2012). Oxytocin modulates cooperation within and competition between groups: an integrative review and research agenda. Horm Behav, 61(3), 419–428. 10.1016/j.yhbeh.2011.12.009

Dumais, K. M., & Veenema, A. H. (2016). Vasopressin and oxytocin receptor systems in the brain: Sex differences and sex-specific regulation of social behavior. Frontiers in Neuroendocrinology, 40, 1–23. 10.1016/j.yfrne.2015.04.003

Faraji, M., Viera-Resto, O. A., Setlow, B., & Bizon, J. L. (2024). Effects of reproductive experience on cost-benefit decision making in female rats [Original Research]. Frontiers in Behavioral Neuroscience, 18.

Ferreira, A. C., & Osório, F. d. L. (2022). Peripheral oxytocin concentrations in psychiatric disorders – A systematic review and methanalysis: Further evidence. Progress in Neuro-Psychopharmacology and Biological Psychiatry, 117, 110561. 10.1016/j.pnpbp.2022.110561

Figueira, R. J., Peabody, M. F., & Lonstein, J. S. (2008). Oxytocin receptor activity in the ventrocaudal periaqueductal gray modulates anxiety-related behavior in postpartum rats. Behav Neurosci, 122(3), 618–628. 10.1037/0735-7044.122.3.618

Fischer-Shofty, M., Shamay-Tsoory, S. G., Harari, H., & Levkovitz, Y. (2010). The effect of intranasal administration of oxytocin on fear recognition. Neuropsychologia, 48(1), 179–184. 10.1016/j.neuropsychologia.2009.09.003

Fuxe, K., Borroto-Escuela, D. O., Romero-Fernandez, W., Ciruela, F., Manger, P., Leo, G., … Agnati, L. F. (2012). On the role of volume transmission and receptor–receptor interactions in social behaviour: Focus on central catecholamine and oxytocin neurons. Brain Research, 1476, 119–131. 10.1016/j.brainres.2012.01.062

Gabriel, D. B. K., Freels, T. G., Setlow, B., & Simon, N. W. (2019). Risky decision-making is associated with impulsive action and sensitivity to first-time nicotine exposure. Behav Brain Res, 359, 579–588. 10.1016/j.bbr.2018.10.008

Goodin, B. R., Ness, T. J., & Robbins, M. T. (2015). Oxytocin - a multifunctional analgesic for chronic deep tissue pain. Curr Pharm Des, 21(7), 906–913. 10.2174/1381612820666141027111843

Gozzi, M., Dashow, E. M., Thurm, A., Swedo, S. E., & Zink, C. F. (2017). Effects of Oxytocin and Vasopressin on Preferential Brain Responses to Negative Social Feedback. Neuropsychopharmacology, 42(7), 1409–1419. 10.1038/npp.2016.248

Hansson, A. C., Koopmann, A., Uhrig, S., Bühler, S., Domi, E., Kiessling, E., … Spanagel, R. (2018). Oxytocin Reduces Alcohol Cue-Reactivity in Alcohol-Dependent Rats and Humans. Neuropsychopharmacology, 43(6), 1235–1246. 10.1038/npp.2017.257

Hodges, T. E., Eltahir, A. M., Patel, S., Bredewold, R., Veenema, A. H., & McCormick, C. M. (2019). Effects of oxytocin receptor antagonism on social function and corticosterone release after adolescent social instability in male rats. Horm Behav, 116, 104579. 10.1016/j.yhbeh.2019.104579

Horn, J. P., & Swanson, L. W. (2012). The autonomic motor system and the hypothalamus. In E. R. Kandel, J. H. Schwartz, T. M. Jessell, S. A. Siegelbaum, & A. J. Hudspeth (Eds.), Principles of neural science (Fifth ed., pp. 2310–2356). McGraw-Hill.

Huber, D., Veinante, P., & Stoop, R. (2005). Vasopressin and Oxytocin Excite Distinct Neuronal Populations in the Central Amygdala. Science, 308(5719), 245–248. 10.1126/science.1105636

Jiang, Y., & Platt, M. L. (2018). Oxytocin and vasopressin increase male-directed threats and vocalizations in female macaques. Scientific Reports, 8(1), 18011. 10.1038/s41598-018-36332-0

Jurek, B., & Meyer, M. (2020). Anxiolytic and Anxiogenic? How the Transcription Factor MEF2 Might Explain the Manifold Behavioral Effects of Oxytocin [Mini Review]. Frontiers in Endocrinology, 11.

Jurek, B., & Neumann, I. D. (2018). The oxytocin receptor: From intracellular signaling to behavior. Physiological Reviews, 98, 1805–1908. 10.1152/physrev.00031.2017

Kapetaniou, G. E., Reinhard, M. A., Christian, P., Jobst, A., Tobler, P. N., Padberg, F., & Soutschek, A. (2021). The role of oxytocin in delay of gratification and flexibility in non-social decision making. Elife, 10. 10.7554/eLife.61844

King, C. E., Gano, A., & Becker, H. C. (2020). The role of oxytocin in alcohol and drug abuse. Brain Res, 1736, 146761. 10.1016/j.brainres.2020.146761

Kirsch, P. (2015). Oxytocin in the socioemotional brain: implications for psychiatric disorders. Dialogues Clin Neurosci, 17(4), 463–476. 10.31887/DCNS.2015.17.4/pkirsch

Klockars, O. A., Klockars, A., Levine, A. S., & Olszewski, P. K. (2018). Oxytocin administration in the basolateral and central nuclei of amygdala moderately suppresses food intake. Neuroreport, 29(6), 504–510. 10.1097/wnr.0000000000001005

Knobloch, H. S., Charlet, A., Hoffmann, L. C., Eliava, M., Khrulev, S., Cetin, A. H., … Grinevich, V. (2012). Evoked axonal oxytocin release in the central amygdala attenuates fear response. Neuron, 73, 553–566. 10.1016/j.neuron.2011.11.030

Kohli, S., King, M. V., Williams, S., Edwards, A., Ballard, T. M., Steward, L. J., … Fone, K. C. F. (2019). Oxytocin attenuates phencyclidine hyperactivity and increases social interaction and nucleus accumben dopamine release in rats. Neuropsychopharmacology, 44(2), 295–305. 10.1038/s41386-018-0171-0

Kosfeld, M., Heinrichs, M., Zak, P. J., Fischbacher, U., & Fehr, E. (2005). Oxytocin increases trust in humans Nature, 435, 673–676. 10.1038/nature03701

Kou, J., Lan, C., Zhang, Y., Wang, Q., Zhou, F., Zhao, Z., … Kendrick, K. M. (2021). In the nose or on the tongue? Contrasting motivational effects of oral and intranasal oxytocin on arousal and reward during social processing. Translational Psychiatry, 11(1), 94. 10.1038/s41398-021-01241-w

Kovács, G. L., Borthaiser, Z., & Telegdy, G. (1985). Oxytocin reduces intravenous heroin self-administration in heroin-tolerant rats Life Sciences, 37(1), 17–26. 10.1016/0024-3205(85)90620-4

Lawson, E. A. (2017). The effects of oxytocin on eating behaviour and metabolism in humans. Nat Rev Endocrinol, 13(12), 700–709. 10.1038/nrendo.2017.115

Lefter, R., Ciobica, A., Antioch, I., Ababei, D. C., Hritcu, L., & Luca, A. C. (2020). Oxytocin Differentiated Effects According to the Administration Route in a Prenatal Valproic Acid-Induced Rat Model of Autism. Medicina (Kaunas), 56(6). 10.3390/medicina56060267

Leong, K.-C., Cox, S., King, C., Becker, H., & Reichel, C. M. (2018). Oxytocin and rodent models of addiction. International Review of Neurobiology, 140, 201–247. 10.1016/bs.irn.2018.07.007

Leslie, M., Leppanen, J., Paloyelis, Y., Nazar, B. P., & Treasure, J. (2019). The influence of oxytocin on risk-taking in the balloon analogue risk task among women with bulimia nervosa and binge eating disorder. Journal of Neuroendocrinology, 31(8). 10.1111/jne.12771

Li, L., Richter, A., & Steinorth, P. (2023). Mental health changes and the willingness to take risks. The Geneva Risk and Insurance Review, 48(1), 31–62. 10.1057/s10713-021-00070-7

Lischke, A., Berger, C., Prehn, K., Heinrichs, M., Herpertz, S. C., & Domes, G. (2012). Intranasal oxytocin enhances emotion recognition from dynamic facial expressions and leaves eye-gaze unaffected. Psychoneuroendocrinology, 37(4), 475–481. 10.1016/j.psyneuen.2011.07.015

Liu, C. M., Hsu, T. M., Suarez, A. N., Subramanian, K. S., Fatemi, R. A., Cortella, A. M., Kanoski, S. E. (2020). Central oxytocin signaling inhibits food reward-motivated behaviors and VTA dopamine responses to food-predictive cues in male rats. Hormones and Behavior, 126, 104855. 10.1016/j.yhbeh.2020.104855

Liu, C. M., Spaulding, M. O., Rea, J. J., Noble, E. E., & Kanoski, S. E. (2021). Oxytocin and Food Intake Control: Neural, Behavioral, and Signaling Mechanisms. International Journal of Molecular Sciences, 22(19).

Liu, Y., Li, S., Lin, W., Li, W., Yan, X., Wang, X., Ma, Y. (2019). Oxytocin modulates social value representations in the amygdala. Nature Neuroscience, 22(4), 633–641. 10.1038/s41593-019-0351-1

Louvel, D., Delvaux, M., Felez, A., Fioramonti, J., Bueno, L., Lazorthes, Y., & Frexinos, J. (1996). Oxytocin increases thresholds of colonic visceral perception in patients with irritable bowel syndrome. Gut, 39(5), 741–747. 10.1136/gut.39.5.741

Love, T. M. (2014). Oxytocin, motivation and the role of dopamine. Pharmacol Biochem Behav, 119, 49–60. 10.1016/j.pbb.2013.06.011

Lu, Q., Lai, J., Du, Y., Huang, T., Prukpitikul, P., Xu, Y., & Hu, S. (2019). Sexual dimorphism of oxytocin and vasopressin in social cognition and behavior. Psychol Res Behav Manag, 12, 337–349. 10.2147/prbm.s192951

Lussier, D., Cruz-Almeida, Y., & Ebner, N. C. (2019). Musculoskeletal Pain and Brain Morphology: Oxytocin’s Potential as a Treatment for Chronic Pain in Aging [Review]. Frontiers in Aging Neuroscience, 11.

MacFadyen, K., Loveless, R., DeLucca, B., Wardley, K., Deogan, S., Thomas, C., & Peris, J. (2016). Peripheral oxytocin administration reduces ethanol consumption in rats. Pharmacology, Biochemistry and Behavior, 140, 27–32. 10.1016/j.pbb.2015.10.014

Marsh, N., Marsh, A. A., Lee, M. R., & Hurlemann, R. (2020). Oxytocin and the Neurobiology of Prosocial Behavior. The Neuroscientist, 27(6), 604–619. 10.1177/1073858420960111

McRae-Clark, A. L., Baker, N. L., Maria, M. M., & Brady, K. T. (2013). Effect of oxytocin on craving and stress response in marijuana-dependent individuals: a pilot study. Psychopharmacology (Berl), 228(4), 623–631. 10.1007/s00213-013-3062-4

Mekhael, A. A., Bent, J. E., Fawcett, J. M., Campbell, T. S., Aguirre-Camacho, A., Farrell, A., & Rash, J. A. (2023). Evaluating the efficacy of oxytocin for pain management: An updated systematic review and meta-analysis of randomized clinical trials and observational studies. Can J Pain, 7(1), 2191114. 10.1080/24740527.2023.2191114

Michalopoulou, P. G., Averbeck, B. B., Kalpakidou, A. K., Evans, S., Bobin, T., Kapur, S., & Shergill, S. S. (2015). The effects of a single dose of oxytocin on working memory in schizophrenia. Schizophr Res, 162(1-3), 62–63. 10.1016/j.schres.2014.12.029

Miller, M. A., Bershad, A., King, A. C., Lee, R., & de Wit, H. (2016). Intranasal oxytocin dampens cue-elicited cigarette craving in daily smokers: a pilot study. Behav Pharmacol, 27(8), 697–703. 10.1097/FBP.0000000000000260

Mitchell, M. R., Weiss, V. G., Beas, B. S., Morgan, D., Bizon, J. L., & Setlow, B. (2014). Adolescent risk taking, cocaine self-administration, and striatal dopamine signaling. Neuropsychopharmacology, 39(4), 955–962. 10.1038/npp.2013.295

Neumann, I. D., & Slattery, D. A. (2016). Oxytocin in General Anxiety and Social Fear: A Translational Approach. Biological Psychiatry, 79(3), 213–221. 10.1016/j.biopsych.2015.06.004

Nishimura, H., Yoshimura, M., Shimizu, M., Sanada, K., Sonoda, S., Nishimura, K., Ueta, Y. (2022). Endogenous oxytocin exerts anti-nociceptive and anti-inflammatory effects in rats. Communications Biology, 5(1), 907. 10.1038/s42003-022-03879-8

Oh, K. S., Kim, E. J., Ha, J. W., Woo, H. Y., Kwon, M. J., Shin, D. W., Lim, S. W. (2018). The Relationship between Plasma Oxytocin Levels and Social Anxiety Symptoms. Psychiatry Investig, 15(11), 1079–1086. 10.30773/pi.2018.08.31

Olivera-Pasilio, V., & Dabrowska, J. (2020). Oxytocin Promotes Accurate Fear Discrimination and Adaptive Defensive Behaviors [Review]. Frontiers in Neuroscience, 14.

Orsini, C. A., Blaes, S. L., Setlow, B., & Simon, N. W. (2019). Recent Updates in Modeling Risky Decision Making in Rodents. In F. H. Kobeissy (Ed.), Psychiatric Disorders: Methods and Protocols (pp. 79–92). Springer New York. 10.1007/978-1-4939-9554-7_5

Orsini, C. A., Hernandez, C. M., Singhal, S., Kelly, K. B., Frazier, C. J., Bizon, J. L., & Setlow, B. (2017). Optogenetic Inhibition Reveals Distinct Roles for Basolateral Amygdala Activity at Discrete Time Points during Risky Decision Making. J Neurosci, 37(48), 11537–11548. 10.1523/JNEUROSCI.2344-17.2017

Orsini, C. A., Trotta, R., T., Bizon, J., L., & Setlow, B. (2015). Dissociable Roles for the Basolateral Amygdala and Orbitofrontal Cortex in Decision-Making under Risk of Punishment. The Journal of Neuroscience, 35(4), 1368. 10.1523/JNEUROSCI.3586-14.2015

Orsini, C. A., Willis, M. L., Gilbert, R. J., Bizon, J. L., & Setlow, B. (2016). Sex differences in a rat model of risky decision making. Behavioral Neuroscience, 130(1), 50–61. 10.1037/bne0000111

Pailing, A. N., & Reniers, R. (2018). Depressive and socially anxious symptoms, psychosocial maturity, and risk perception: Associations with risk-taking behaviour. PLoS One, 13(8), e0202423. 10.1371/journal.pone.0202423

Patel, N., Grillon, C., Pavletic, N., Rosen, D., Pine, D. S., & Ernst, M. (2015). Oxytocin and vasopressin modulate risk-taking. Physiol Behav, 139, 254–260. 10.1016/j.physbeh.2014.11.018

Pedersen, C. A., Ascher, J. A., Monroe, Y. L., & Prange, A. J. (1982). Oxytocin induces maternal behavior in virgin female rats. Science, 216(4546), 648–650. 10.1126/science.7071605

Pedersen, C. A., & Boccia, M. L. (2003). Oxytocin antagonism alters rat dams’ oral grooming and upright posturing over pups. Physiol Behav, 80(2-3), 233–241. 10.1016/j.physbeh.2003.07.011

Peters, S., Slattery, D. A., Uschold-Schmidt, N., Reber, S. O., & Neumann, I. D. (2014). Dose-dependent effects of chronic central infusion of oxytocin on anxiety, oxytocin receptor binding and stress-related parameters in mice. Psychoneuroendocrinology, 42, 225–236. 10.1016/j.psyneuen.2014.01.021

Pincus, D., Kose, S., Arana, A., Johnson, K., Morgan, P., Borckardt, J., Nahas, Z. (2010). Inverse Effects of Oxytocin on Attributing Mental Activity to Others in Depressed and Healthy Subjects: A Double-Blind Placebo Controlled fMRI Study [Original Research]. Frontiers in Psychiatry, 1.

Plessow, F., Marengi, D. A., Perry, S. K., & Lawson, E. A. (2021). Oxytocin Administration Increases Proactive Control in Men with Overweight or Obesity: A Randomized, Double-Blind, Placebo-Controlled Crossover Study. Obesity (Silver Spring), 29(1), 56–61. 10.1002/oby.23010

Pujara, M., & Koenigs, M. (2014). Mechanisms of reward circuit dysfunction in psychiatric illness: prefrontal-striatal interactions. Neuroscientist, 20(1), 82–95. 10.1177/1073858413499407

Qi, J., Yang, J.-Y., Wang, F., Zhao, Y.-N., Song, M., & Wu, C.-F. (2009). Effects of oxytocin on methamphetamine-induced conditioned place preference and the possible role of glutamatergic neurotransmission in the medial prefrontal cortex of mice in reinstatement. Neuropharmacology, 56, 856–865. 10.1016/j.neuropharm.2009.01.010

Qiu, F., Qiu, C. Y., Cai, H., Liu, T. T., Qu, Z. W., Yang, Z., Hu, W. P. (2014). Oxytocin inhibits the activity of acid-sensing ion channels through the vasopressin, V1A receptor in primary sensory neurons. Br J Pharmacol, 171(12), 3065–3076. 10.1111/bph.12635

Rae, M., Lemos Duarte, M., Gomes, I., Camarini, R., & Devi, L. A. (2022). Oxytocin and vasopressin: Signalling, behavioural modulation and potential therapeutic effects. British Journal of Pharmacology, 179(8), 1544–1564. 10.1111/bph.15481

Reddy, L. F., Lee, J., Davis, M. C., Altshuler, L., Glahn, D. C., Miklowitz, D. J., & Green, M. F. (2014). Impulsivity and risk taking in bipolar disorder and schizophrenia. Neuropsychopharmacology, 39(2), 456–463. 10.1038/npp.2013.218

Rhianne, S., Tylah Ms, D., Erin, L., Nicholas, E., & Michael, B. (2023). A Novel Clinical-stage Small Molecule Targeting the ‘Dark Side’ of Opioid Addiction. Journal of Pharmacology and Experimental Therapeutics, 385(S3), 168. 10.1124/jpet.122.262620

Rickenbacher, E., Perry, R. E., Sullivan, R. M., & Moita, M. A. (2017). Freezing suppression by oxytocin in central amygdala allows alternate defensive behaviours and mother-pup interactions. Elife, 6. 10.7554/eLife.24080

Schorscher-Petcu, A., Sotocinal, S., Ciura, S., Dupré, A., Ritchie, J., Sorge, R. E., … Mogil, J. S. (2010). Oxytocin-induced analgesia and scratching are mediated by the vasopressin-1A receptor in the mouse. J Neurosci, 30(24), 8274–8284. 10.1523/jneurosci.1594-10.2010

Simon, N. W., Gilbert, R. J., Mayse, J. D., Bizon, J. L., & Setlow, B. (2009). Balancing Risk and Reward: A Rat Model of Risky Decision Making. Neuropsychopharmacology, 34(10), 2208–2217. 10.1038/npp.2009.48

Simon, N. W., Montgomery, K. S., Beas, B. S., Mitchell, M. R., LaSarge, C. L., Mendez, I. A., … Setlow, B. (2011). Dopaminergic modulation of risky decision-making. J Neurosci, 31(48), 17460–17470. 10.1523/jneurosci.3772-11.2011

Smith, A. S., Korgan, A. C., & Young, W. S. (2019). Oxytocin delivered nasally or intraperitoneally reaches the brain and plasma of normal and oxytocin knockout mice. Pharmacol Res, 146, 104324. 10.1016/j.phrs.2019.104324

Smith, C. J. W., Poehlmann, M. L., Li, S., Ratnaseelan, A. M., Bredewold, R., & Veenema, A. H. (2017). Age and sex differences in oxytocin and vasopressin V1a receptor binding densities in the rat brain: focus on the social decision-making network. Brain Structure and Function, 222(2), 981–1006. 10.1007/s00429-016-1260-7

Song, Z., & Albers, H. E. (2018). Cross-talk among oxytocin and arginine-vasopressin receptors: Relevance for basic and clinical studies of the brain and periphery. Front Neuroendocrinol, 51, 14–24. 10.1016/j.yfrne.2017.10.004

Spetter, M. S., Feld, G. B., Thienel, M., Preissl, H., Hege, M. A., & Hallschmid, M. (2018). Oxytocin curbs calorie intake via food-specific increases in the activity of brain areas that process reward and establish cognitive control. Scientific Reports, 8(1), 2736. 10.1038/s41598-018-20963-4

St Onge, J. R., & Floresco, S. B. (2009). Dopaminergic modulation of risk-based decision making. Neuropsychopharmacology, 34(3), 681–697. 10.1038/npp.2008.121

Stevens, D. (2005). Understanding our fears. Nature Reviews Neuroscience, 6(6), 420–420. 10.1038/nrn1694

Strawbridge, R. J., Ward, J., Cullen, B., Tunbridge, E. M., Hartz, S., Bierut, L., Smith, D. J. (2018). Genome-wide analysis of self-reported risk-taking behaviour and cross-disorder genetic correlations in the UK Biobank cohort. Translational Psychiatry, 8(1), 39. 10.1038/s41398-017-0079-1

Takayanagi, Y., & Onaka, T. (2021). Roles of Oxytocin in Stress Responses, Allostasis and Resilience. Int J Mol Sci, 23(1). 10.3390/ijms23010150

Tapp, D. N., Singstock, M. D., Gottliebson, M., S., & McMurray, M. S. (2020). Central but not peripheral oxytocin administration reduces risk-based decision-making in male rats. Hormones and Behavior, 125(104840). 10.1016/j.yhbeh.2020.104840

Terburg, D., Scheggia, D., Triana Del Rio, R., Klumpers, F., Ciobanu, A. C., Morgan, B., van Honk, J. (2018). The Basolateral Amygdala Is Essential for Rapid Escape: A Human and Rodent Study. Cell, 175(3), 723–735.e716. 10.1016/j.cell.2018.09.028

Tunstall, B. J., Kirson, D., Zallar, L. J., McConnell, S. A., Vendruscolo, J. C. M., Ho, C. P., … Vendruscolo, L. F. (2019). Oxytocin blocks enhanced motivation for alcohol in alcohol dependence and blocks alcohol effects on GABAergic transmission in the central amygdala. PLoS Biol, 17(4), e2006421. 10.1371/journal.pbio.2006421

Uvnäs-Moberg, K. (2023). The physiology and pharmacology of oxytocin in labor and in the peripartum period. American Journal of Obstetrics and Gynecology. 10.1016/j.ajog.2023.04.011

Valstad, M., Alvares, G. A., Andreassen, O. A., Westlye, L. T., & Quintana, D. S. (2016). The relationship between central and peripheral oxytocin concentrations: a systematic review and meta-analysis protocol. Systematic Reviews, 5(1), 49. 10.1186/s13643-016-0225-5

Viviani, D., & Stoop, R. (2008). Opposite effects of oxytocin and vasopressin on the emotional expression of the fear response. In I. D. Neumann & R. Landgraf (Eds.), Progress in Brain Research (Vol. 170, pp. 207–218). Elsevier. 10.1016/S0079-6123(08)00418-4

Walter, M. H., Abele, H., & Plappert, C. F. (2021). The role of oxytocin and the effect of stress during childbirth: Neurobiological basics and implications for mother and child. Frontiers in Endocrinology, 12(742236). 10.3389/fendo.2021.742236

Wang, D., Yan, X., Li, M., & Ma, Y. (2017). Neural substrates underlying the effects of oxytocin: a quantitative meta-analysis of pharmaco-imaging studies. Social Cognitive and Affective Neuroscience, 12(10), 1565–1573. 10.1093/scan/nsx085

Wang, T., Shi, C., Li, X., Zhang, P., Liu, B., Wang, H., … Xu, Z. D. (2018). Injection of oxytocin into paraventricular nucleus reverses depressive-like behaviors in the postpartum depression rat model. Behav Brain Res, 336, 236–243. 10.1016/j.bbr.2017.09.012

Weber, R. A., Logan, C. N., Leong, K.-C., Peris, J., Knackstedt, L., & Reichel, C. M. (2018). Regionally specific effects of oxytocin on reinstatement of cocaine seeking in male and female rats. International Journal of Neuropsychopharmacology, 21(7), 677–686. 10.1093/ijnp/pyy025

Wheeler, A. R., Truckenbrod, L. M., Cooper, E. M., Betzhold, S. M., Setlow, B., & Orsini, C. A. (2023). Effects of fentanyl self-administration on risk-taking behavior in male rats. Psychopharmacology (Berl), 240(12), 2529–2544. 10.1007/s00213-023-06447-y

Whitton, A. E., Treadway, M. T., & Pizzagalli, D. A. (2015). Reward processing dysfunction in major depression, bipolar disorder and schizophrenia. Curr Opin Psychiatry, 28(1), 7–12. 10.1097/yco.0000000000000122

Williams, J. R., Insel, T. R., Harbaugh, C. R., & Carter, C. S. (1994). Oxytocin administered centrally facilitates formation of a partner preference in female prairie voles (Microtus ochrogaster). J Neuroendocrinol, 6(3), 247–250. 10.1111/j.1365-2826.1994.tb00579.x

Xin, Q., Bai, B., & Liu, W. (2017). The analgesic effects of oxytocin in the peripheral and central nervous system. Neurochemistry International, 103, 57–64. 10.1016/j.neuint.2016.12.021

Yao, S., & Kendrick, K. M. (2022). Effects of Intranasal Administration of Oxytocin and Vasopressin on Social Cognition and Potential Routes and Mechanisms of Action. Pharmaceutics, 14(2). 10.3390/pharmaceutics14020323

Yoshida, M., Takayanagi, Y., Inoue, K., Kimura, T., Young, L. J., Onaka, T., & Nishimori, K. (2009). Evidence that oxytocin exerts anxiolytic effects via oxytocin receptor expressed in serotonergic neurons in mice. J Neurosci, 29(7), 2259–2271. 10.1523/jneurosci.5593-08.2009

Zebhauser, P. T., Macchia, A., Gold, E., Salcedo, S., Burum, B., Alonso-Alonso, M., Brem, A.-K. (2022). Intranasal oxytocin modulates decision-making depending on outcome predictability—A randomized within-subject controlled trial in healthy males. Biosciences, 10(12). 10.3390/biomedicines10123230

Zhou, L., Sun, W.-L., Young, A. B., Lee, K., McGinty, J. F., & See, R. E. (2014). Oxytocin reduces cocaine seeking and reverses chronic cocaine-induced changes in glutamate receptor function. International Journal of Neuropsychopharmacology, 18(1), 1–11. 10.1093/ijnp/pyu009

